# Optimizing viral genome subsampling by genetic diversity and temporal distribution (TARDiS) for Phylogenetics

**DOI:** 10.1101/2021.01.15.426832

**Authors:** Simone Marini, Carla Mavian, Alberto Riva, Marco Salemi, Brittany Rife Magalis

## Abstract

TARDiS for Philogenetics is a novel tool for optimal genetic sub-sampling. It optimizes both genetic diversity and temporal distribution through a genetic algorithm. TARDiS, along with example data sets and a user manual, is available at https://github.com/smarini/tardis-phylogenetics

## 1 Introduction

Viral genetic sequence data can be used for integrated phylogenetic and population genetic, or phylodynamic, analysis to trace viral evolutionary patterns, as well as spatiotemporal origin and dissemination of viral and bacterial pathogens [5]. Phylodynamic tools, such as NextStrain [6] and BEAST [23, 1], are now routinely utilized to monitor evolution and population dynamics of epidemics based on real-time deposition of pathogen sequences in databases (e.g. GenBank, HIVdatabases, GISAID) [22, 14, 12, 10, 15, 13, 24]. Not unlike traditional epidemiological analysis, however, these methods can significantly be affected by sampling bias [7], and sampling during outbreaks are rarely performed randomly from a representative, stratified population [21]. Not only do the quality and quantity of sequences vary per country, but even regional sample collection policies tend to be inconsistent over time, as exemplified by the inherent sampling bias of SARS-CoV-2 strains, collected through convenience sampling, and sequenced during the early pandemic phase [15]. Moreover, continuous generation of new sequences can very quickly approach information overload. For example, as of January 15th, 2021, ∼375,000 sequences have been deposited in GISAID (SARS-CoV-2) database, with a number of countries either over or under represented compared of their actual infection prevalence. In such cases, full dataset analyses cannot be accomplished, as computational tools are not designed to handle hundred of thousands of sequences. In order to reduce computational complexity, subsampling must often be performed [8], typically using an approach that maximizes genetic diversity among subpopulations [2] (e.g., countries or regions [8]), which increases phylogenetic signal in the dataset, thus improving phylodynamic inference over convenience sampling. Besides enhancing signal for statistical phylogenetic inference, reliable estimates of significant events in the context of space and time also require sufficient temporal signal in the dataset [7], or distribution of sampling over time, to calibrate reliable molecular clocks [19]. Despite sampling strategies pose a significant threat to conclusions drawn from phylodynamic inference, this problem has received so far insufficient attention [4]. Hall *et al*. (2016) [7] were able to demonstrate that sampling sequences uniformly with respect to both space and time leads to better solutions than optimizing sampling solely by genomic diversity. There currently exists no tool to aid researchers to optimize pathogens’ sequences sub-sampling with respect to space, time, and genetic diversity. In what follows, we introduce TARDiS (Temporal And diveRsity Distribution Sampler), a machine learning approach designed to optimize phylogenetic subsampling according to both genetic diversity and temporal distribution for user-defined subpopulations.

## 2 Methods

### 2.1 Genetic Algorithm

TARDiS implements a genetic algorithm (GA) [9, 3] optimizing genetic diversity and time sampling distributions criteria for any set of viral or bacterial genomes. The output consists of user-defined number *n* of optimally subsampled genomes from a complete dataset of *N* genomes. Briefly, the algorithm is initialized as a population of random individuals. Each individual is a solution to the problem, i.e., a subsample of size *n* genomes. Each individual is characterized by a fitness score, reflecting how well that particular individual (solution) performs on the given problem. In our case, fitness is measured as a combination of genetic diversity (i.e., how diverse are the genomes represented by the individual), and time distribution (i.e., how evenly distributed are the genomes represented by the individual along the epidemic timeline).

#### Genetic diversity maximization

We aim to recover a subsample of genomes as genetically diverse as possible. To do so, we first need to calculate the genetic distance between all possible genome pairs, represented by a square distance matrix *D*, with *N* rows and columns. The user can provide their own distance matrix, or let TARDiS compute it using the Jukes-Cantor substitution model. We calculate the genetic diversity fitness *F*_*gd*_ of a subset as

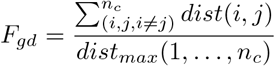

where *n*_*c*_ are all the genome pairs (*i, j*) ∈ *n*, with *i* ≠ *j*; *dist*(*i, j*) is the genetic distance of a genome pair (*i, j*); and *dist*_*max*_(1, …, *n*_*c*_) is the sum of the genetic diversities of the maximum *n* elements of the distance matrix *D*, representing a theoretical upper bound to force fitness ∈ [0, 1], with a higher value representing a better *F*_*gd*_.

#### Time distribution optimization

Our objective is to recover a subsample of *n* genomes that are evenly distributed along the considered time interval. Intuitively, if *n* = 10 and the time interval is 10 days, we would like to consider one genome per day. We can thus calculate the ideal time distribution *I*_*td*_ as a date vector of *n* elements, starting with the first available date *d*_*f*_, ending with the last available date *d*_*l*_, and having the remaining *n* − 2 elements distanced with a 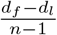 interval. The worst possible time distribution *W*_*td*_, on the other hand, is a time distribution concentrated into a single specific date (i.e., all samples collected on the same day). We measure the time distribution fitness *F*_*td*_ as

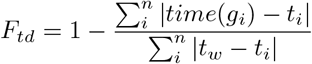

where *time*(*g*_*i*_) is the collection date of the i-th genome, *t*_*i*_ is i-th date in *I*_*td*_, and *t*_*w*_ is i-th date in *W*_*td*_. In other words, *F*_*td*_ is bounded ∈ [0, 1], with a higher value representing a better *F*_*td*_. The final fitness *F* of a specific individual is calculated as

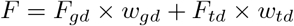

where *w*_*gd*_ and *w*_*td*_ are user-defined weights to set the importance of genetic diversity and time distribution, respectively.

#### GA operators

Once a population is generated, fitness is calculated for each individual. Individuals are then chosen and combined to produce a new population, in an iterative fashion. To generate a novel individual, TARDiS uses three operators: selection, mutation, and crossover. The selection operator is based on deterministic tournament selection with *k* = 5 [3]. Briefly, two sets of *k* individuals are randomly chosen, and the individual with the highest fitness is selected from each set. The crossover operator combines two tournament winners *A* and *B* into a new individual *C* by keeping the all the *g* genomes ∈ (*A* ∪ *B*), and randomly selecting 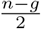 genomes ∈ (*A* ∩ *B*) − (*A* ∪ *B*). To help avoid local maxima, each newly generated individual *C* has a 0.08 probability of mutating [9, 3]. A mutation is defined as swapping a genome of individual *C* with one randomly chosen from the remaining, non-(*A*|*B*) genome pool. Note also that the user defines a fraction of the population that is randomly created (and thus not evolved) for each generation. Another user-defined value determines elitism, i.e., the fraction of best genomes (ranked by fitness) to be copied without modifications in the next generation.

### 2.2 Case study: subsampling a rising epidemic

We simulated a growing epidemic using a stochastic, agent-based model [11] with limited migration between ten subpopulations, or regions (a, …, j). Details on simulation paramaters can be found in the supplementary materials. We ran TARDiS on a single simulated dataset, subsampling 40 genomes per region (with the exception of region f, with 37 genomes available) for 50 generations, with a population of 1000 individuals per generation, of which 85% were evolved, 10% were newly generated, and 5% were elite. The phytools package [20] in R [17] was used for joint likelihood reconstruction of discrete ancestral states [16] according to subpopulation for each internal node of the subsampled trees. Transition rates among discrete states along tree nodes were considered to be equal *a priori*. Migration rates between states were then re-estimated, as described in the supplementary methods. We compared the results obtained both with (*w*_*g*_*d* = 1, *w*_*t*_*d* = 1) and without considering time distribution (*w*_*g*_*d* = 1, *w*_*t*_*d* = 0). Our simulation indicated that better results are obtained by considering time distribution: the overall migration rate root mean squared error (RMSE) decreased by 17% (0.035 to 0.029). Eight representative clades were then chosen for which the majority of taxa consisted of a single sub-population and were consistent across true and subsampled trees (Figure 1, A). The RMSE decreased by 43.4% (from 25.37 considering only genetic diversity (t0) to 14.36 if we include time distribution (t1). The addition of a temporal weighting component for an exponentially growing population can act to both increase and decrease representation of earlier time points (e.g., weeks 15 and 12, respectively; Figure 1, B). However, representation of week 1 of the epidemic was increased from 0% to 5%. As the early stages of an epidemic, and time nearing the root of the tree, represent periods of high epidemiological and phylogenetic uncertainty, sample representation during this time is critical for reliable phylodynamic inference and thus contributed to the loss of error in our estimates.

**Figure 1:**
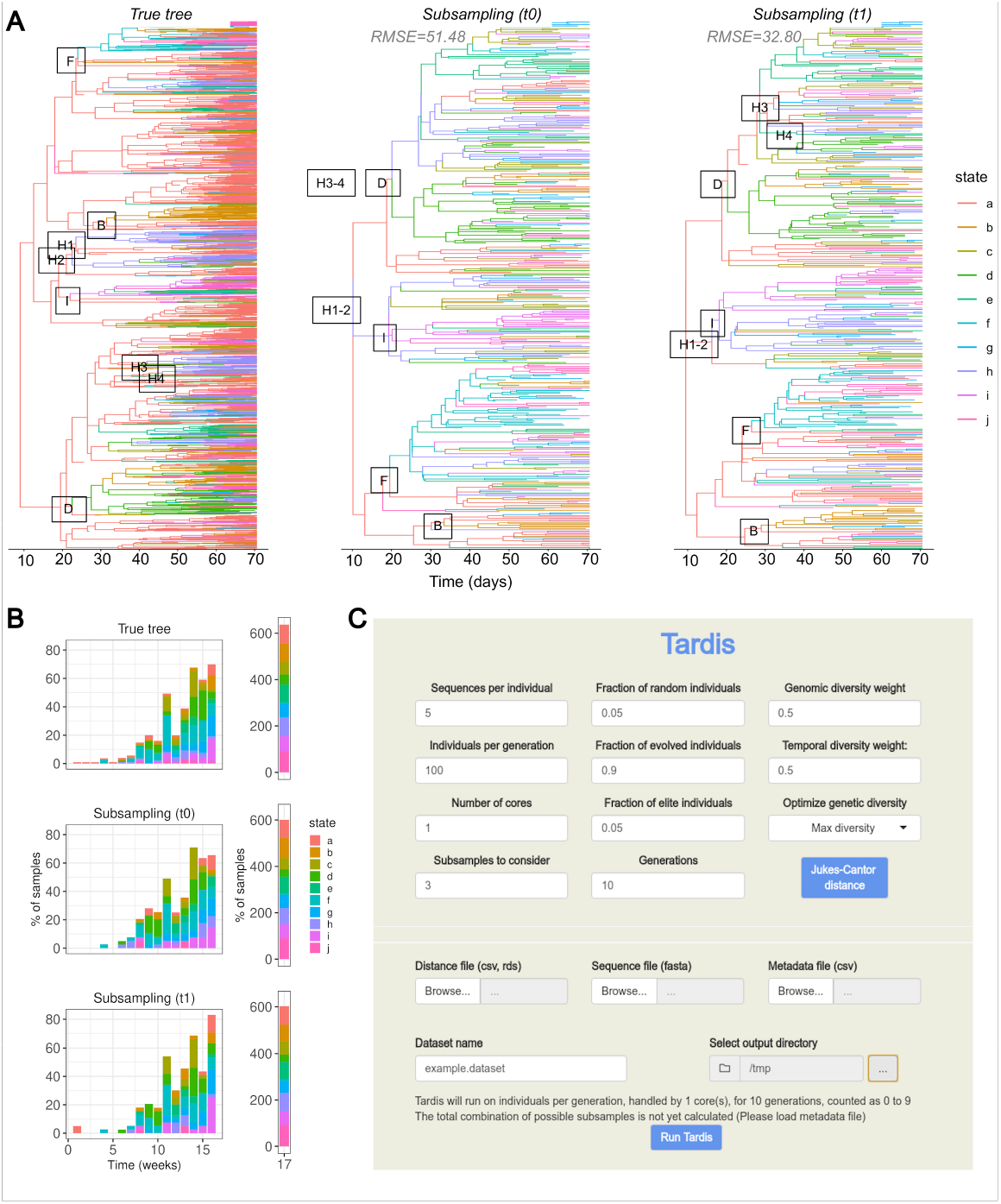
A) True and subsampled trees with representative clades. Eight representative clades were chosen for which the majority of taxa consisted of a single subpopulation, or state, and were consistent across true and subsampled trees. Root mean squared error (RMSE) was calculated for the true times to the most recent common ancestors (TMRCAs) and estimated TMRCAs across the five representative clades for the subsampled tree with (t1) and without consideration of time (t0). (B) Temporal distribution of samples per subpopulation for true and subsampled trees with (t1) and without consideration of time (t0). (C) Screenshot of the TARDiS graphical user interface.

#### Simulation principles

Each subpopulation was allowed to emerge from the initially infected population (a) with a mean probability of [initial] infection of 0.02 (standard deviation [sd] of 0.005). Each infected individual within a subpopulation was then allowed to migrate to another subpopulation with a mean probability of 0.01 (sd=0.005). The number of contacts for each individual was picked from a normal distribution with a mean of 4 (sd=2). The probability of transmission (when a contact occurs) was provided in the form of a threshold function: prior to 5 days (sd=3), the host was not able to transmit, but after that time, the individual was able transmit with a mean probability of 0.05 (sd=0.005), representing an incubation period for the simulated virus.

Each infected individual was removed from the simulation (representing death, recovery, etc.) after 14 days. The described parameters resulted in a basic reproductive number (*R*_0_) of approximately 1.6 for the epidemic. The simulated epidemic was run for 365 days or until a total of 10,000 hosts were infected. For each of the ten subpopulations, individuals belonging to that subpopulation were binned according to week of removal (i.e., seven-day intervals) and subsam-pled according to an exponential distribution (rate=5), representing idealistic sampling of a population proportional to the size of the epidemic and resulting in a range [37, 844] of sampled individuals for each subpopulation (state). The original transmission tree was pruned, leaving only the remaining sampled individuals.

A molecular clock, or constant evolutionary rate across all branches of the tree, was assumed, allowing branches separating noes within the tree to be scaled in both time and genetic distance. Nucleotide (A,C,G,T) sequences were thus simulated along the tree using a general time reversible evolutionary model, with rate matrix (0.32512,1.07402,0.26711,0.25277,2.89976,1.00000) and nucleotide frequencies (0.299,0.183,0.196,0.322). A gamma distributed of rate variation across sites (alpha=2.35) was also used, with a proportion (0.60) of sites considered to be invariable. Branch lengths were scaled by a factor of 8e-04 (representing an approximate evolutionary rate in substitutions/site/year). Sequence simulation was performed in Seq-Gen [18].

Migration rates for each of the ten subpopulations were calculated as a function of the number of transitions between subpopulation states (non-reversible) along each branch within the tree and the frequency (*F*) of the initial subpopulation among tree tips. I.e, for *w* branches with transitions between subpopulations, and *x* branches with specifically transitions from *i* (node at earlier time point) to *j* (node at more recent time point),

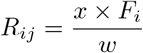

## 3 Implementation

TARDiS is implemented as a command-line tool based on NextFlow, suitable for analyzing large datasets in an HPC environment, and as a GUI based on R/Shiny for ease-of-use and experimentation 1. TARDiS is available at https://github.com/smarini/tardis-phylogenetics.

